# Neutral models of *de novo* gene emergence suggest that gene evolution has a preferred trajectory

**DOI:** 10.1101/2023.02.05.527172

**Authors:** Bharat Ravi Iyengar, Erich Bornberg-Bauer

## Abstract

New protein coding genes can emerge from genomic regions that previously did not contain any genes, via a process called *de novo* gene emergence. To synthesize a protein, DNA must be transcribed as well as translated. Both processes need certain DNA sequence features. Stable transcription requires promoters and a polydenylation signal, while translation requires at least an open reading frame (ORF). We develop mathematical models based on mutation probabilities, and the assumption of neutral evolution, to find out how quickly genes emerge and are lost. We also investigate the effect of the order by which DNA features evolve, and if sequence composition is biased by mutation rate. We rationalize how genes are lost much more rapidly than they emerge, and how genes with long ORFs preferentially arise in regions that are already transcribed. Our study not only answers some fundamental questions on the topic of *de novo* emergence but also provides a modeling framework for future studies.

## 1 Introduction

Organisms evolve new traits by expanding their functional genome. Evolution of new genes is one of the ways by which new traits can emerge. The definition of a gene is complicated, and has been changing constantly (Gerstein *et al*., 2007). We use the following working definition of a gene: a gene is a region of the genome that gives rise to a functional product, that is an RNA or a protein. The next big challenge is to define what a function is, and how to classify gene functions. One of the easiest definition of a gene product’s function is its ability to improve the survival of the organism under one or more environmental conditions (Keeling *et al*., 2019). Proteins perform the most diverse kinds of molecular functions ranging from catalysis of biochemical reactions to formation of cellular structures (Berg *et al*., 2002), whereas RNAs that do not encode proteins like mRNAs, or participate in protein synthesis like rRNAs and tRNAs, are mostly involved in regulation of gene expression (Iyengar *et al*., 2014; Statello *et al*., 2021).

In this study we focus on the evolution of genes that encode new proteins. New protein coding genes frequently arise via duplication of existing protein coding genes. These duplicated genes then genetically and functionally diverge from their parent genes by accumulating mutations (Long *et al*., 2003; Nüsvall *et al*., 2012). Recently it has been shown that new protein coding genes can arise independent of gene duplication, from genomic sequences that did not previously encode any protein. This phenomenon is called *de novo* gene emergence (Tautz and Domazet-Lošo, 2011; Zhao *et al*., 2014; Schmitz and Bornberg-Bauer, 2017; Vakirlis *et al*., 2017; Van Oss and Carvunis, 2019). *De novo* protein coding genes can emerge from intergenic sequences, non-coding RNA genes, introns, and even regions that partially overlap with existing protein coding genes (Tautz and Domazet-Lošo, 2011; Zhao *et al*., 2014; Vakirlis *et al*., 2017; Van Oss and Carvunis, 2019; Prabh and Rödelsperger, 2019). To express a protein, a genomic sequence should be transcribed as well as translated. Therefore, the gene needs sequence features that enable both these processes.

The primary requirement for transcription is recruitment of RNA polymerase at the gene’s promoter (Lenhard *et al*., 2012). A part of the promoter, called the core promoter is the region that determines the start of transcription (Haberle and Stark, 2018). The second requirement for a productive transcription is its termination at appropriate sites. Even though pervasive transcription is widespread in eukaryotic genomes, most RNAs are quickly eliminated by RNA degrading enzymes (Schmid and Jensen, 2018). Most eukaryotic mRNAs and long non-coding RNAs, are polyadenylated at their 3’ termini, which makes them resistant to degradation. Polyadenylation, which also marks the end of transcription (Richard and Manley, 2009), is facilitated by sequences known as polyadenylation signals (Proudfoot, 2011). In prokaryotes, the end of transcription is determined by sequences known as terminators (Santangelo and Artsimovitch, 2011). Thus the synthesis of a stable RNA requires promoters, as well as termination or polyadenylation signal sequences.

The second major requirement for protein expression is translation of the transcribed mRNA. The most fundamental requirement for translation is the presence of an open reading frame (ORF). Usually, an mRNA needs additional features to initiate protein synthesis. These features include ribosome binding sites in prokaryotes (Omotajo *et al*., 2015), and Kozak consensus sequences in eukaryotes (Kozak, 1986; Acevedo *et al*., 2018; Noderer *et al*., 2014).

Although gain of transcription and translation features guarantees *de novo* birth of a protein coding gene, it does not ensure that the gene would persist in the genome for many generations. The newly born gene can lose the features as easily as it gained them unless it has been fixed in the genome, for example via evolutionary selection. Specifically, if the protein synthesized by the *de novo* gene provides a fitness advantage to the host organism, it will undergo positive selection (Keeling *et al*., 2019). Simultaneously, the protein should have low toxicity and cost of synthesis, to survive purifying selection. A common mechanism by which protein mediated toxicity occurs is by misfolding and aggregation of proteins (Bucciantini *et al*., 2002; Hartl, 2017).

In this study we develop probabilistic models of *de novo* gene emergence. We focus on *de novo* genes emerging from non-genic sequences, also known as “proto-genes” (Carvunis *et al*., 2012; Van Oss and Carvunis, 2019). We model *de novo* gene emergence as a two step process. In the first step, a non-genic DNA sequence gains transcriptional features that allows it to express an untranslated (non-coding) RNA. In the second step this non-coding RNA gene acquires translational features that allows it to express a protein. Alternatively, translational features can emerge in the non-genic DNA before transcriptional features. Specifically, we calculate the probabilities by which promoters, polyadenylation signals and ORFs emerge, and are lost, based on the rates at which different kinds of DNA mutations naturally occur (mutation bias). With these probabilities we estimate whether transcriptional and translational features preferentially evolve in a specific order. Using a similar approach, we calculate how random mutations affect protein composition. In our models, we assume that the proteins expressed from these proto-genes provide no fitness advantage and are not toxic to the host organism. Thus our models are based on neutral evolution. Our models predict that there is indeed a preferred sequence of DNA feature evolution during *de novo* emergence, even under the assumption of neutrality.

## 2 Results

In this work, we developed mathematical models to estimate the rates and probabilities of *de novo* gene emergence, as well as gene loss. A proto-gene emerges from non-genic DNA, when the latter mutates to gain sequence features necessary for transcription and translation (Carvunis *et al*., 2012; Van Oss and Carvunis, 2019). Both transcription and translation are complex processes involving many biomolecular complexes that work in concert. Here, we focus on the minimal requirements for these processes to occur. Specifically for transcription, we require a gene to have one of the two core promoter motifs, the TATA-box or the Initiator element (Inr), and a polyadenylation (poly-A) signal sequence. We chose these two promoter motifs because they are the most abundant type of core promoters across diverse eukaryotes (Lenhard *et al*., 2012; Haberle and Stark, 2018). We defined the poly-A signal as a transcription requirement because it is crucial for transcript stability and its export to cytoplasm (Richard and Manley, 2009; Stewart, 2019). For translation, we only require the gene to have an open reading frame (ORF). Furthermore, we assume that the ORF is uninterrupted by introns.

Using simple probability models, we calculate the likelihood of finding these features in a DNA sequence by random chance, and the rate at which these features are gained and lost due to random mutations. These probability models are essentially described by two kinds of probability. First is the probability of finding a DNA sequence feature (such as TATA box or an ORF; Section 4.2.1). This probability depends on the nucleotide distribution given by the GC-content. The second kind of probability is the transition probability, that is the probability of a DNA sequence mutating to another (Section 4.2.2). Transition probabilities that describe the gain and loss of DNA sequence features, depend on mutation rate, mutation bias, and GC-content. Our model of DNA sequence evolution is in principle similar to a theoretical model described in a previous study (Behrens and Vingron, 2010). In this study, the authors have calculated the time required for different transcription factor binding sites to emerge in gene promoters. We extend this principle to calculate not only the probability of sequence emergence but also the likelihood that an emerged sequence will remain intact or be lost (Section 4.3).

For all calculations we used a default GC-content of 0.42 which is reasonably close to the total genomic GC-content of human (0.41, Merchant *et al*., 2007), *Drosophila melanogaster* (0.416, Gramates *et al*., 2022), and *Saccharomyces cerevisiae* (0.38, Wood *et al*., 2002). We used a mutation rate of 7.8 × 10^−9^ mutations per nucleotide position per generation, which corresponds to spontaneous mutation rate estimated in a *Drosophila melanogaster* population that was subjected to periodic population bottlenecks (mutation accumulation line, Schrider *et al*., 2013). We used mutation bias data (Table 1) from the same study (Schrider *et al*., 2013) which was in agreement with another previous study on human pseudogenes (Zhang and Gerstein, 2003).

**Table 1:** Mutation bias probabilities for different nucleotide mutations. `A:T` denotes an `A`-`T` base pair in a double stranded DNA. Thus `A → G` mutation on one DNA strand would cause a `T → C` mutation on the complementary strand. We describe the other mutations in the same way.

| Substitution | Probability($\mu$) |
| --- | --- |
| A:T→T:A | 0.056 |
| A:T→G:C | 0.243 |
| A:T→C:G | 0.074 |
| G:C→A:T | 0.483 |
| G:C→T:A | 0.075 |
| G:C→C:G | 0.069 |

Using a similar approach, we predict how the nucleotide distribution and random mutations shape protein composition and evolution. Overall, our models define a null hypothesis under which protein coding DNA sequences evolve neutrally through mutation pressure alone.

### 2.1 How likely is gene loss?

If a gene is found in one species but not in sister taxa, then the most common assumption is that this gene emerged only in one specific species. We asked what is the chance that the gene was present in a common ancestor but was lost in all lineages except one. To this end we calculated the probability of gene emergence and gene loss (Section 4.3). Specifically, the probability of gene gain can be defined as the sum of three probabilities. First is the probability that an ORF already exists and is not lost due to mutations (*P*_*ORF-stay*_, Section 4.4), while a mutation causes transcription to emerge (*P*_*RNA-gain*_, Section 4.5). Second is the probability that an ORF emerges (*P*_*ORF-gain*_, Section 4.4) in a region of DNA that is already transcribed and continues to be transcribed (*P*_*RNA-stay*_, Section 4.5). Third is the probability that neither of the two features already exist and both emerge at the same time due to mutations (this probability is very small and is negligible). We note that the probability of transcription gain is dependent on the existence of an ORF, such that it is higher when the ORF is already present. This dependence exists because an ORF does not have stop codons (`TAA`, `TAG` and `TGA`) in its sequence. This in turn, restricts the number of positions where a poly-A signal, that has a `TAA` in its sequence, can exist (Section 4.6). Because of the same reason, the probability of ORF gain is higher when the DNA is already transcribed. Therefore, we additionally calculated the conditional probabilities of transcription gain (*P*_*RNA-gain ORF*_) and ORF gain (*P*_*ORF-gain RNA*_) given the condition that an ORF and a transcript already exists, respectively. We thus define the total probability of gene gain 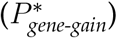 as:

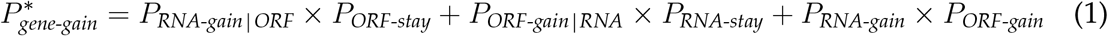

Equation 1 describes the total probability of gene gain that includes the probability that the gene was not previously present. To more precisely address the question of whether a gene that was absent in an ancestor but emerges in a descendant, we calculated the corresponding conditional probability of gene gain:

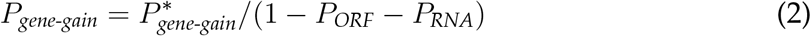

We note that the difference between the total probability 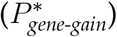 and the conditional probability (*P*_*gene-gain*_) of gene gain, is very small.

Next, we calculated the probability of gene loss, given the gene (transcription and ORF) is already present(*P*_*gene-loss*_). It is the sum of the probabilities that transcription (*P*_*RNA-loss*_, Section 4.5) or the ORF is lost due to mutations (*P*_*ORF-loss*_, Section 4.4). Specifically:

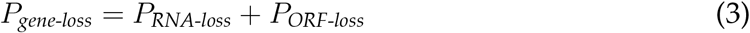

The probability that a gene is lost *n* times independently, is (*P*_*gene-loss*_)^*n*^. To find out how many independent gene loss events are as likely as a single gene gain event, we calculate the ratio of logarithms (log-log ratio) of gene gain and gene loss for protogenes with an ORF ranging from 30 – 300 codons in length.

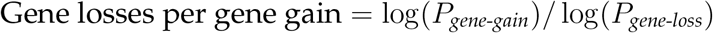

For example, a log-log ratio value of 2 would indicate that two independent gene loss events are as likely as a single gene gain event. We calculated this ratio for genes with different ORF lengths, because ORF gain and loss probabilities depend on the length of the ORF. We found that for ORF longer than 30 codons, at least two independent gene loss events can occur in the time frame of one gene gain event (Figure 1A). To understand how GC-content determines the likelihood of gene gain relative to gene loss, we calculated the log-log ratio of the two probability values for proto-genes with different GC-content (Figure 1B), and that contain ORFs of different lengths (codons). We found that the effect of GC-content on the log-log ratio depends on the length of the ORF. Specifically, the log-log ratio steadily increases with GC-content ranging from 0.3 to 0.6, when the proto-gene contains an ORF with 30 codons. When the ORF has 60 – 90 codons, the log-log ratio initially decreases with GC content and then increases. For ORFs containing more than 120 codons, the log-log ratio decreases with increasing GC-content (within the analysed range). Overall, gene loss could occur at least two times independently in the time frame of one gene gain event, irrespective of the GC content.

**Figure 1:**
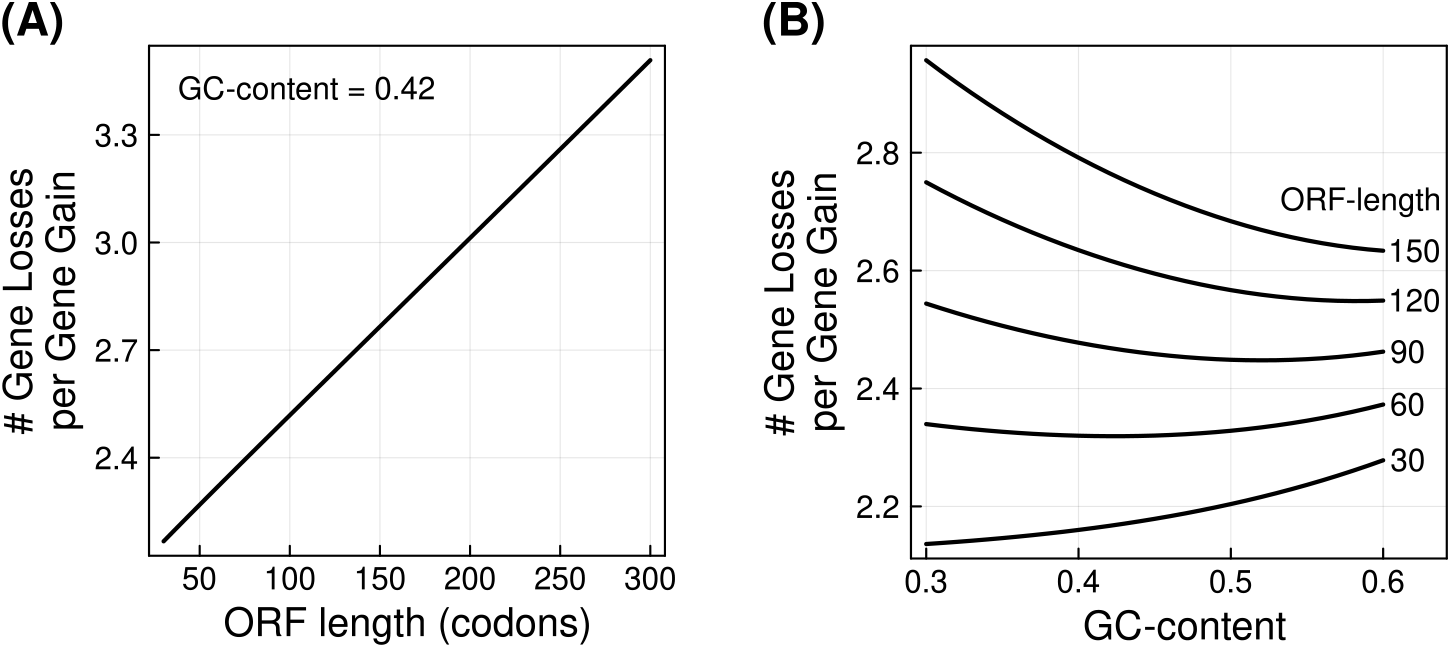
Genes are more likely to be independently lost twice, than being born once. The vertical axis in both panels shows the number of independent gene loss events (Equation 3) that can occur relative to one gene gain event (Equation 2) – log(*P*_*gene-gain*_)*/* log(*P*_*gene-loss*_). The horizontal axis shows the number of codons in the ORF **(A)**, and the GC-content **(B)**.

We performed an analogous analysis on just the ORFs (and not the whole genes). That is, we calculated the ratio of logarithms of ORF gain and ORF loss. We found that for ORFs longer than 142 codons, the likelihood of two independent ORF loss events is more than or equal to a single ORF gain event.

Overall, our analysis suggests that a proto-gene expressed in only one species may not necessarily mean that it emerged for the first time in this species.

### 2.2 Does *de novo* gene emergence follow a preferred trajectory of events?

For a proto-gene to emerge from a non-genic DNA sequence, both transcription and ORF need to emerge. That is, probability of gene emergence is equal to the product of probabilities of transcription gain and ORF gain. Thus it may appear that the order of the occurrence of these two events does not matter. However, gene emergence is much more likely when one of the two features already exists (Equation 1) and therefore it has two possible trajectories – ORF emerges first (ORF-first) or transcription emerges first (RNA-first). To understand whether *de novo* gene emergence has a preferred trajectory, we calculated two probabilities. First, the probability that ORF exists and mutations cause gain of transcription but no disruption of the ORF. This probability denotes the trajectory where ORF emerges first (*P*_*ORF-first*_, Equation 4).

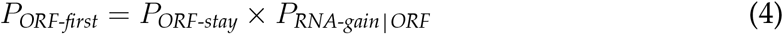

The second probability (*P*_*RNA-first*_, Equation 5) that denotes the trajectory where transcription emerges first, is the probability that DNA is already transcribed and mutations cause a gain of ORF but do not eliminate transcription.

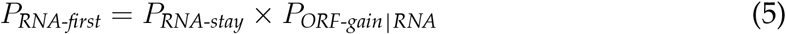

We calculated the log-transformed ratio of *P*_*ORF-first*_ and *P*_*RNA-first*_, such that a positive value of the ratio would mean that transcription gain for an untranscribed ORF (ORF-first) is more likely than ORF gain in an already transcribed DNA (RNA-first). In other words, the ORF-first hypothesis is more feasible. Likewise, a negative value of the ratio would suggest that the RNA-first hypothesis is more feasible. We found that protogenes with an ORF containing 60 codons or more, preferentially emerge RNA-first, whereas proto-genes with shorter ORFs emerge ORF-first (Figure 2A). More specifically, the likelihood of the ORF-first trajectory relative to that of the RNA-first trajectory decreases with increasing ORF length u a minimum value, and then increases. To understand if GC-content determines the evolutionary trajectory of *de novo* emergence, we calculated the probabilities of the ORF-first and the RNA-first trajectory for different values of GC-content. We found that proto-genes with short ORF can emerge ORF-first at high GC-content. For example, the ORF-first trajectory is more likely at a GC-content of 0.6, for ORFs with 30 – 36 codons (Figure 2B).

**Figure 2:**
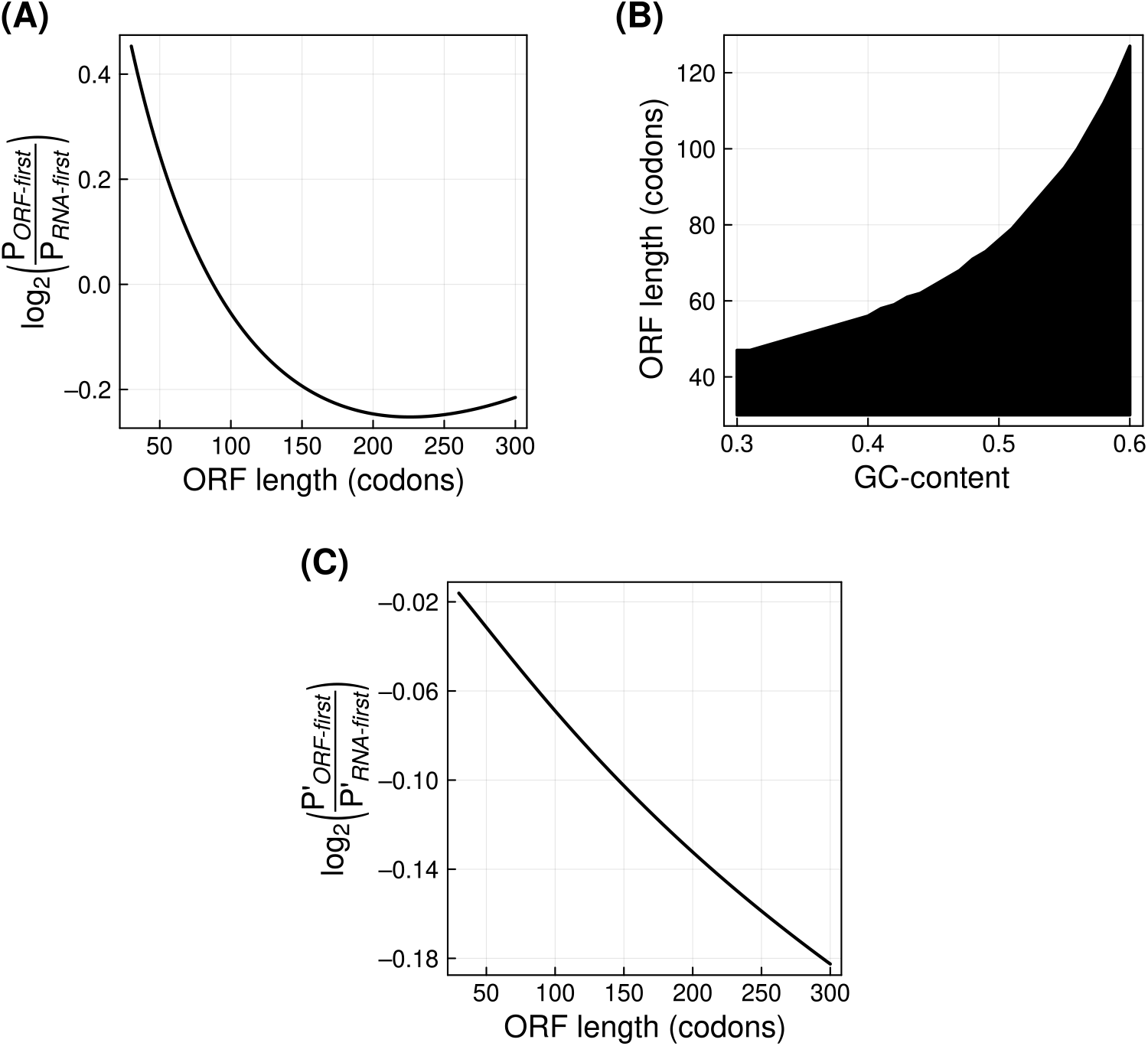
Proto-genes with ORFs longer than 59 codons, emerge RNA-first, whereas protogens with shorter ORFs emerge ORF-first. The vertical axis shows the log_2_ transformed ratio of the probabilities of the ORF-first and the RNA-first trajectories, described as **(A)** a single-step process (Equations 4 & 5) and **(C)** a two-step process (Equations 6 & 7). A negative value suggests that RNA-first trajectory is more likely than ORF-first trajectory, and *vice versa*. In both panels, the horizontal axis denotes the number of codons in the ORF. **(B)** Filled area denotes the ORF length in codons (vertical axis) and the GC-content values (horizontal axis) at which ORF-first trajectory is more likely than RNA-first trajectory..

A more stringent definition of the ORF-first hypothesis would describe a two-step process. In the first step, an ORF emerges in an untranscribed region of DNA but transcription does not emerge. In the second step transcription emerges, while ORF stays intact (Equation 6). Likewise, a RNA-first hypothesis can be defined by a two-step probability. In the first step transcription emerges in a non-genic DNA (that does not have an ORF), and in the second step an ORF emerges, while transcription remains intact (Equation 7)

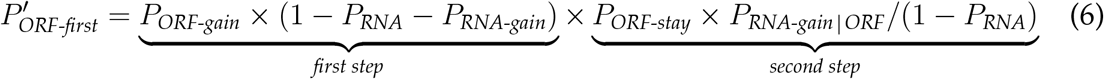

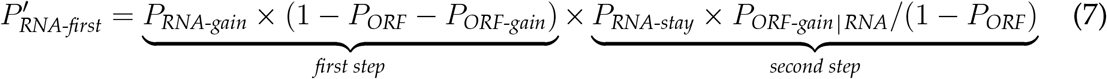

With this stringent definition, proto-genes with every analysed ORF length (30 – 300) preferentially emerge via the RNA-first trajectory (Figure 2C). Furthermore, the twostep RNA-first trajectory becomes increasingly more probable than the two-step ORF-first trajectory, with increasing ORF length.

Overall, long ORFs are likely to be lost before transcription emerges, and hence they preferentially emerge in existing transcripts. Conversely, short ORFs are more likely to stay undisrupted until the emergence of transcription.

### 2.3 Would extensive transcription loss suggest negative selection of toxic proteins?

Proto-genes that do not provide any fitness benefit to an organism may not be fixed in populations via natural selection. These genes may be lost due to mutation pressure. Some newborn proto-genes can also encode toxic proteins, that may aggregate or interfere with physiology in some other way. These genes would thus be eliminated from the population genomes via negative selection. We note that an ORF is lost if the start codon is mutated, the stop codon is mutated to an amino acid encoding codon (non-stop mutation), or if an amino acid encoding codon is mutated to a stop codon (premature stop/non-sense mutation). However, it is likely that a non-stop mutation or a non-sense mutation, can still result in translation of a protein (extended or truncated, respectively). Furthermore, non-stop mutations can also lead to cellular toxicity (Choe *et al*., 2016). Thus ORF loss does not ensure elimination of toxic proteins, which in turn suggests that transcription loss may more effectively inactivate the associated genes.

To understand which is the most probable mechanism of gene loss, we compared the probabilities of ORF loss (*P*_*ORF-loss*_) and transcription loss (*P*_*RNA-loss*_). We found that ORF loss is more probable than transcription loss, especially so when the ORFs are long (Figure 3A). At low GC-content (0.3 – 0.34) ORF loss is less likely than RNA-loss, for short ORFs (30 codons; Figure 3B).

**Figure 3:**
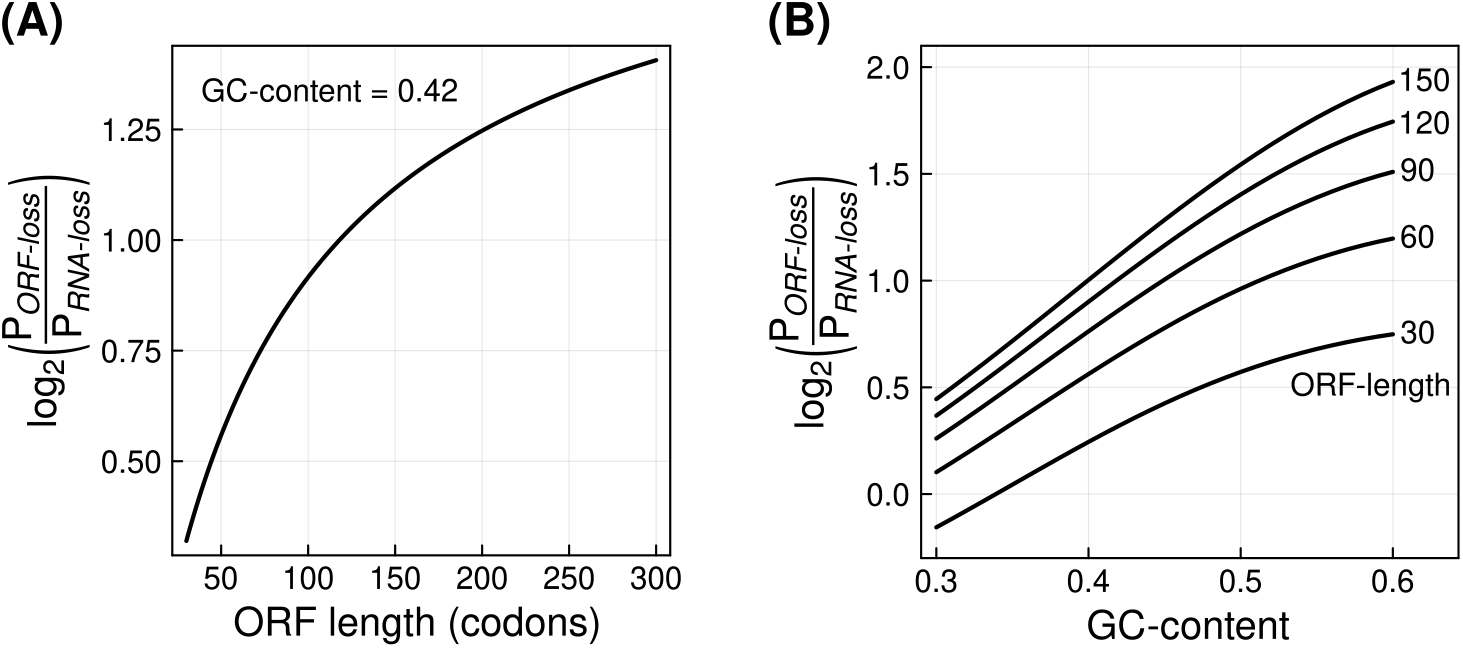
ORF loss in proto-genes is more probable than transcription loss. The horizontal axis denotes the number of codons in the ORF **(A)** and GC-content **(B)**. The vertical axis shows the log_2_ transformed ratio of ORF loss and RNA loss probabilities, such that a positive value means ORF loss is more probable than RNA loss, and *vice versa*.

We made our analysis of gene loss more stringent by comparing the probabilities of two evolutionary scenarios, both of which begin with an intact proto-gene. In the first scenario, the ORF remains intact whereas the transcription is lost. The probability of this scenario (*P*_*onlyORF-loss*_) is described by:

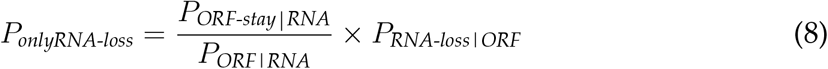

We note that the probability that an ORF is not lost due to mutations (*P*_*ORF-stay RNA*_), includes the probability that the ORF already exists (*P*_*ORF RNA*_). To describe the conditional probability of ORF remaining undisrupted, given that it already exists, we divide *P*_*ORF-stay RNA*_ by *P*_*ORF RNA*_.In the second scenario, transcription stays intact while ORF is lost. The probability of this scenario is:

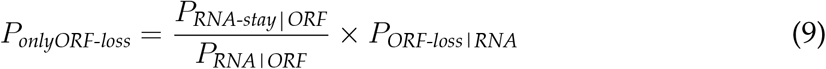

We find that the second scenario (only ORF loss) is more plausible for all the analysed ORFs lengths. Furthermore, the ratio of *P*_*onlyORF-loss*_ and *P*_*onlyRNA-loss*_ has nearly identical values and trend across different ORF lengths (data not shown), as the ratio of *P*_*ORF-loss*_ and *P*_*RNA-loss*_ shown in Figure 3.

Thus, widespread transcription loss can suggest a negative selection on protein expression.

### 2.4 Does mutation bias shape protein composition?

In previous sections, we showed that mutation bias affects the rate of *de novo* gene emergence and loss. We next turned our attention to whether this bias affects the composition (and thereby the chemistry) of proteins encoded by proto-genes. To this end, we first asked if the expected frequency of different amino acids is uniform, and found that it is not uniform. Amino acids like leucine (L) and serine (S) have a higher expected frequency than other amino acids. On the other hand, amino acids like methionine (M) and tryptophan (W) are less probable (Figure 4A). This non-uniformity is to a great extent determined by number of degenerate codons for an amino acid. However, GC-content also plays a role. For example, leucine, arginine and serine, all have six codons each. However, leucine is more likely to be encoded than serine in a random stretch of DNA, given a uniform GC-content of 0.42 (Figure 4A). Overall, our analysis is roughly in agreement with many previous studies (Ohta and Kimura, 1971; Shen *et al*., 2006; Gardini *et al*., 2016). However, notable differences exist between our analysis and that of the previous studies because the latter primarily focused on characterized proteins and not random proto-gene derived protein sequences.

**Figure 4:**
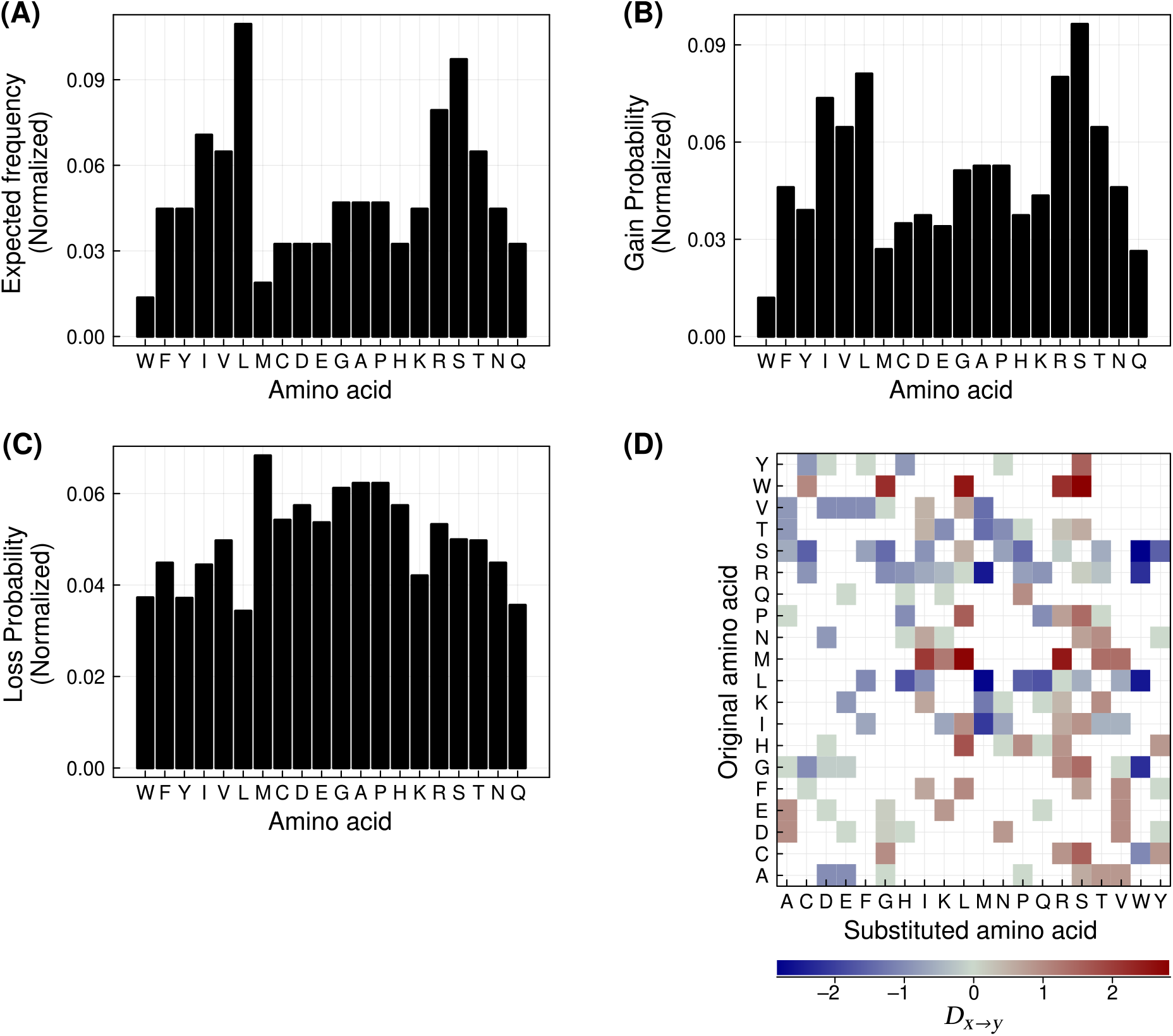
Mutation bias and GC-content shape protein composition. Panels **(A), (B)** and **(C)** show expected frequency, gain probability (Equation 10), and loss probability (Equation 11) for different amino acids (horizontal axes). The vertical axis in all these three subplots show the normalized values of these measures such that their sum over all amino acids is equal to one. Panel **(D)** shows the directionality (colored tiles, *D*_*x→y*_, Equation 12) of the substitution from an amino acid (vertical axis) to another (horizontal axis). Positive values of *D*_*x→y*_ are shown in shades of red, whereas negative values are shown in shades of blue. We only show substitutions that can be realized by a single nucleotide mutation but we include all possible nucleotide mutations in the directionality calculations for these substitutions. In all panels, amino acids are ordered based on their chemical similarity (Kim *et al*., 2009).

Next, we aimed to find out if some amino acid substitutions are more probable than the others. To this end, we calculated the gain probability (Equation 10) for every amino acid. Specifically for every amino acid *x*, we calculated the average probability that it substitutes any of the other nineteen amino acids, due to mutations. More precisely, the total probability that an amino acid *y* will mutate to an amino acid *x* is the product of the expected frequency of amino acid *y* (*P*_*y*_), and the substitution rate from *x* to *y* (*µ*_*y→x*_). The gain probability for an amino acid *x* is then defined as:

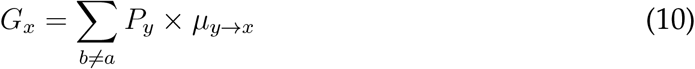

We found that amino acid gain probabilities were also non-uniform across different amino acids (Figure 4B) but they had a strong positive correlation with the expected frequency (Pearson *ρ* = 0.955, *P <* 10^−10^).

Next, we calculated how easily is an amino acid mutated to any other amino acid. Specifically, the conditional loss probability for an amino acid (*L*_*x*_), given the amino acid already exists, is defined as:

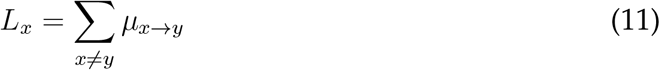

The loss probability was also not uniformly distributed (Figure 4C) but it did not significantly correlate with the expected frequency of the amino acids (Pearson *ρ* = −0.255, *P* = 0.2818).

Next, we investigated if some amino acid substitutions are more common than the others. As expected, amino acid substitutions that require more than one nucleotide change were much less probable than those needing just a single nucleotide change. Therefore, we focused our analysis on amino acid substitutions whose total probabilities were more than that of a double nucleotide mutation (note that this procedure identifies amino acid substitutions that are possible via single nucleotide mutations but it does not exclude multinucleotide mutations from the calculation of amino acid substitution probabilities). Although single nucleotide mutations are more probable than double mutations, the different single mutations do not occur at the same rate due to mutation rate bias. Thus an amino acid substitution (for example K→E) may not be as likely to occur as the reverse substitution (E→K). That is substitutions between a pair of amino acids may have a directionality. Although many previous studies have explored the likelihood of different amino acid substitutions (Ohta and Kimura, 1971; Dayhoff *et al*., 1978; Henikoff and Henikoff, 1992; Gonnet *et al*., 1992; Jones *et al*., 1992; Whelan and Goldman, 2001; Schneider *et al*., 2005; Kosiol *et al*., 2007; Le and Gascuel, 2008), none to our knowledge have focused primarily on the directionality of these substitutions. To verify if such a directionality exists, we calculated the log transformed ratio of forward and reverse substitution probability, *D*_*x→y*_, for every pair of amino acids, *x* and *y*. A positive value of *D*_*x→y*_ means that *x* to *y* substitution is more likely than *y* to *x* substitution, and *vice versa*.

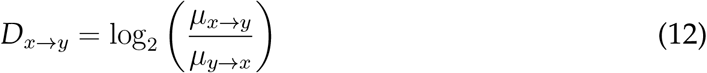

As we mentioned in the previous paragraph, we restricted this analysis to substitutions that can be achieved via single nucleotide mutations. We found that most substitutions indeed had a preferred direction (*D*_*xy*_ ≠ 0, Figure 4D). The median absolute valueof *D*_*xy*_ was 0.905 and the maximum value was 2.836. That is, in 38 out of 75 amino acid pairs we analysed, one direction of substitution was at least 1.87 times more probable than the other. For example, E→K substitution is 1.8 times more probable than K→E substitution. This may appear as a small number but many such unidirectional substitutions occurring in an evolving protein could indicate a directional selection. We also performed an analogous analysis where we defined the substitution directionality using the ratio of corresponding amino acid gain probabilities (Equation 10). Using this definition reduces the range of directionality values (*D*_*xy*_) such that its median value is 0.38. Although this more stringent definition reduces the intensity of the directionality, it does not change the qualitative results.

A previous study found that mutations that cause hydrophobic amino acids to appear on protein surfaces, can lead to evolution of protein dimers (Hochberg *et al*., 2020). The authors also suggested that such an evolutionary process may be widespread because mutation bias tends to facilitate emergence of hydrophobic amino acids. They based this argument on another study that estimated mutation rate bias in bacteria (Hershberg and Petrov, 2010). The authors (Hochberg *et al*., 2020) further suggested that hydrophobic amino acids are more frequently found in random protein sequences.

We asked if our model also makes similar predictions. To this end, we first calculated the percentage of amino acids in a protein sequence that are expected to be hydrophobic. A popular hydrophobicity scale of amino acids is based on solubility of an amino acid in water or ethanol (Kyte and Doolittle, 1982). However, this scale does not classify tryptophan as a hydrophobic amino acid. Another hydrophobicity scale estimated by a different study (Wimley and White, 1996) is more biologically realistic and is based on rates of transfer of different amino acids between water and a hydrophobic medium. This study analysed two hydrophobic media – a lipid bilayer and octanol, and also considered the effect of peptide bonds in the calculations. Therefore we used the hydrophobicity scale by Wimley and White (1996) for classifying amino acids such that hydrophobic amino acids have a negative free energy change of transfer from water to hydrophobic media. Conversely, a hydrophilic amino acid would have a positive value of the free energy change. Based the octanol hydrophobicity scale, we classified the following amino acids as hydrophobic – cysteine (C), phenylalanine (F), isoleucine (I), leucine (L), methionine (M), valine (V), tryptophan (W) and tyrosine (Y). With the lipid bilayer hydrophobicity scale, valine is not classified as hydrophobic, possibly because it does not integrate well in the bilayer. Therefore we used the octanol scale for our calculations.

We found that *∼*37.5% of amino acids in a protein sequence are expected to be hydrophobic. This finding is broadly in agreement with the previous study that places this percentage at 40% (Hochberg *et al*., 2020). We note that cytosolic proteins may require more than 42% of their constituent amino acids to be hydrophobic, in order to fold efficiently (Dill, 1985). Next, we asked if a protein sequence in general tends to be hydrophobic. Specifically, we calculated expected hydrophobicity 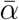, Equation 13) of a protein sequence, based on expected frequency of different amino acids (*P*_*x*_, Figure 4) and their hydrophobicity values (*α*_*x*_).

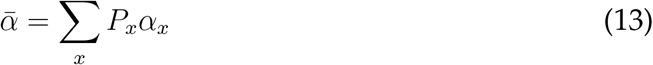

We found that the expected hydropathy is higher than zero (0.39), which suggests that random protein sequences are not on an average, hydrophobic.

To find out if random mutations indeed cause hydrobphobic amino acids to occur in protein sequences, we calculated the probability that any mutation substitutes a nonhydrophobic amino acid with a hydrophobic amino acid (*P*_*α-gain*_, Equation 14). Likewise, we calculated the probability that any mutation causes a hydrophobic to non-hydrophobic amino acid substitution (*P*_*α-loss*_, Equation 15).

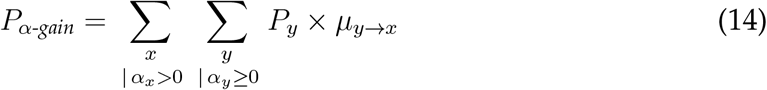

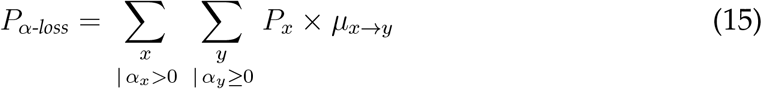

Next, we calculated the ratio of *P*_*α-gain*_ and *P*_*α-loss*_, and found that it is 1.114, which suggests that gain of a hydrophobic amino acid is slightly more probable than its loss. We performed a complementary analysis where we calculated the average change in hydrophobicity due to any random mutation 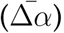, defined by:

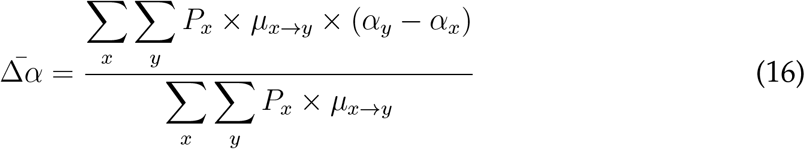

We found the average change in hydrophobicity to be −0.033, which suggests that on an average mutations cause a very small increase in hydrophobicity. We reiterate that hydrophobic amino acids have a negative value hydrophobicity and thus a negative change in hydrophobicity also denotes a shift towards a more hydrophobic protein. To focus exclusively on non-hydrophobic to hydrophobic amino acid substitutions, we excluded amino acid pairs where both amino acids are either hydrophobic or are non-hydrophobic. That is, we only analysed pairs of amino acids that have opposite signs of hydrophobicity. With this focused analysis, we find that the average change in hydrophobicity is even smaller (−3.7 × 10^−4^).

Overall our analyses show that random mutations may preferentially cause hydrophobic amino acids to accumulate in protein sequence but this effect is very small.

## 3 Discussion

In this work, we addressed some fundamental questions about *de novo* emergence of protogenes. Broadly, we asked how quickly these genes can emerge and be lost, whether their birth and death has a preferred trajectory of mutational events, and how the composition of their protein sequence is determined. To this end, we developed mathematical models and used them to address specific questions. These models are based on mutation probability and mutation rate bias, and represent neutral evolution where DNA sequences are not evolving under natural selection. Thus our models provide an opportunity to test the alternative hypothesis that some proto-genes are evolving under selection.

We show that genes can be lost much more rapidly than they are gained, such that a gene can be independently lost two to three times in the time required for it to emerge. In a hypothetical case, a proto-gene may be present in a species (A) but is absent in two sister taxa (B and C). Two evolutionary scenarios could explain this observation. In the first scenario, the proto-gene simply emerged for the first time in species A. The second scenario posits that the proto-gene was present in the common ancestor of the three species but was lost in both species B and species C. Our models suggest that the second scenario is more likely. Future phylogenetic studies on proto-genes should consider this finding for making inferences on the dynamics of gene gain and loss.

Within the limitations of our model assumptions, we answer the long standing question of which trajectory of mutational events leads to *de novo* emergence. That is, whether ORF emerges first or transcription emerges first (Schmitz and Bornberg-Bauer, 2017). We show that, in the absence of selection, long ORFs (*≥*60 codons) are likely to emerge in existing transcripts (transcription emerges first), whereas short ORFs are likely to emerge before transcription emerges in the genomic locus. However, if both transcription and ORF are absent in a genomic locus, then *de novo* emergence is more likely to occur transcription-first. Long non-coding RNAs may be a potential source of proto-genes with long ORFs but this evolutionary process may be constrained by a different set of factors. A previous study has shown that new transcripts frequently arise in regions overlapping existing genes. These new transcripts can harbor ORFs which in turn leads to *de novo* gene emergence (Blevins *et al*., 2021). This is an example of RNA-first trajectory. However, the ORFs present in these genes may be under indirect selection pressure that acts on the overlapping gene sequences. Therefore, they may be less frequently mutated or lost.

Although our model is based on the assumption of neutral evolution, one can speculate the effects of positive selection on the proto-gene. For example, if the gene produces a protein that is beneficial to the organism, then selection would cause the fixation of the gene as soon as the ORF emerges via RNA-first trajectory. On the other hand, the selection would have no effect on an ORF that is not transcribed. This is based on the assumption that the ORF sequence has no other function (for example, regulation), or is not under indirect selection (for example, if it overlaps with an existing gene). Thus positive selection will intensify the preference for RNA-first trajectory.

We show that transcription loss is less likely than the loss of long ORFs. However, it may be difficult to find direct evidence for such a phenomenon. Specifically, it is difficult to infer from data if transcription was lost or never emerged. If the protein encoded by an ORF is indeed toxic, it would be silenced or purged rapidly in microevolutionary time scales. One way of inferring negative selection from transcription loss may involve correlation of ORF sequence variation and transcription status. Mutations within an ORF sequence are much more probable than mutations that disrupt transcription or an ORF. This is so because the entire ORF sequence can have a higher number of mutable sites than a promoter. If an ORF variant is untranscribed in many populations (species or taxa), it is possibly toxic.

A large volume of work exists on the likelihood of amino acid substitutions and many of these models are widely used in phylogenetics studies (Ohta and Kimura, 1971; Dayhoff *et al*., 1978; Henikoff and Henikoff, 1992; Gonnet *et al*., 1992; Jones *et al*., 1992; Whelan and Goldman, 2001; Schneider *et al*., 2005; Kosiol *et al*., 2007; Le and Gascuel, 2008). However, commonly used amino acid substitution models such as PAM and BLOSUM (Dayhoff *et al*., 1978; Henikoff and Henikoff, 1992) do not focus on directionality of mutations (the substitution matrices are symmetric). Our aim was to highlight how proteins can evolve under mutation pressure and that mutation bias causes some substitutions to occur more frequently than the others. Furthermore, we do not base our substitution probability calculations on known protein sequences but rather on random protein sequences that are more likely to to be encoded in proto-genes. Using our models, we show that for a pair of amino acids, one direction of substitution can be significantly more likely than the other. If a substitution occurs more frequently than it is expected to occur in our neutral model, then directional evolution can be a possible explanation for the observation. We also show that random mutations can lead to a small increase in a protein’s hydrophobicity. Hydrophobic amino acids can facilitate folding (Dill, 1985), but can also cause proteins to aggregate (Hochberg *et al*., 2020). Thus the effect of hydrophobicity increasing mutations on a protein’s stability and toxicity, is dependent on where they occur in the protein structure.

Like all theoretical models, our models are not without limitations. First, we assume that nucleotides are uniformly distributed according to GC-content. This is not true as nucleotide composition can vary significantly across the genome. Mutation rates are also not uniform across the genome and can vary significantly (Monroe *et al*., 2022). However, our models can be adapted to investigate specific genomic loci, where information on the local GC-content and mutation rate can more accurately determine the *de novo* gene evolution in these loci. Second, our models are based on parameters such as mutation rate and mutation bias, which we obtained from published data on human and *D*.*melanogaster* genomes (Zhang and Gerstein, 2003; Schrider *et al*., 2013). The accuracy of the models’ prediction will depend on the accuracy of parameters. However, mutation rate itself only determines the values of different probabilities. Our results, which are based on probability ratios, will not significantly change with a different value of mutation rate. More importantly, different species may have a different spectrum of mutations (Cano *et al*., 2022), especially for species that are highly distant from each other. Thus our inferences based on our chosen parameters will not universally apply to all species. Third, we assume that all mutations are independent. This is not the case for tandem mutations, where multinucleotide mutations occur more frequently than their expected frequency under the assumption of mutational independence (Harris and Nielsen, 2014). Fourth, we did not model the effect of many regulatory sequence features such as enhancers, transcription factor binding sites, 5’ and 3’ untranslated regions, and Kozak consensus sequence, that can influence the evolution of proto-genes. For example *de novo* emergence is very likely to occur near enhancers (Majic and Payne, 2020). We instead focused on minimal requirements for transcription and translation. However, our modeling framework allows inclusion of any number of sequence features in the calculations as long as their sequences can be defined (Section 4.3). Finally, we ignore insertions, deletions and transpositions as mechanisms of mutation. Insertions and deletions (indels) are reported to be less frequent than substitutions (Schrider *et al*., 2013), and both indels and transpositions cause larger sequence alterations than point mutations. Our modeling methodology also allows incorporation of these mechanisms as a source of mutations.

A famous aphorism about theoretical models says “all models are wrong but some are useful” (Box, 1979). Our models are not an exception. They are not 100% accurate but they are useful for making predictions that can lead to a more focused experimental validation. These models can be made more complex to accommodate diverse molecular mechanisms driving gene evolution. Therefore our work opens up an opportunity for theoretical and computational biologists to develop models using our modeling framework, that more accurately describe their system of interest.

## 4 Materials and Methods

### 4.1 Mutation probabilities

We calculated nucleotide substitution probabilities based on mutation rate and mutation rate bias data. Specifically, we used a mutation rate (*λ*) of 7.8 × 10^−9^ mutations per nucleotide position per generation (Schrider *et al*., 2013). We derived our mutation bias parameters from two published studies, the first on *Drosophila melanogaster* (Schrider *et al*., 2013), and the second on humans (Zhang and Gerstein, 2003). Table 1 shows the exact values of mutation bias probabilities that we used in this study.

### 4.2 Probabilities of finding, gaining, and losing a DNA sequence

#### 4.2.1 Probability of finding a DNA sequence

We calculated the probability of finding a DNA sequence based on global nucleotide frequency distributions, given by the GC-content. We used a GC-content of 0.42 for our calculations. Specifically, the probability of finding either a `G` or a `C` is: *S* = 0.5×GC-content. The probability of finding an `A` or a `T` is: *W* = 0.5 *− S*. Using these values, we calculated the probability of finding a DNA sequence motif by chance. For example, the probability of finding the sequence `ATG` would be: *W × W × S*.

#### 4.2.2 Probability of gaining a DNA sequence

We calculated the probability of gaining a DNA sequence due to mutations using GC-content, mutation rate and mutation bias. Specifically, we calculated the probability that a DNA sequence does not exist, and it emerges due to specific nucleotide mutations. More precisely, this probability is the product of two other probabilities. The first is the probability of finding a DNA sequence (*x*) that is not the sequence of interest. The second probability is that this sequence *x* mutates to the sequence of interest. To explain this calculation better, we use the example of `CTA` mutating to `ATG`. The first probability, that is the probability of finding `CTA` by chance is *SW* ^2^. `CTA` mutates to `ATG` via two nucleotide mutations (`C`→`A` and `A`→`G`). Thus the probability of this DNA change would be the probability of two nucleotide mutations (*λ*^2^) multiplied by two mutation bias probabilities (`G:C`→`T:A` and `A:T`→`G:C`). Overall, the chance of `CTA` mutating to `ATG` would be:

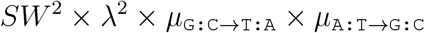

Next, we calculated the probability that every nucleotide triplet that is not `ATG`, mutates to `ATG`. This can happen via one, two, or three nucleotide mutations. The sum of all these mutation probabilities is the probability of `ATG` gain.

Using the same principle we calculated the gain probability of any DNA sequence motif (of any length or composition). We excluded insertions and deletions as a mechanism of gain of small DNA sequences that we analysed in this study.

#### 4.2.3 Probability of losing a DNA sequence

We calculated the loss of a DNA sequence motif using the same principle we used for calculating gain probabilities. However, we defined loss probability as a conditional probability, that is we assume that the DNA sequence of interest already exists in the genome. For example, the loss probability of a specific `ATG` sequence would be the sum of probabilities of `ATG` mutating to any of the other 63 nucleotide triplets (via one, two, or three nucleotide mutations). We use conditional loss probabilities by default, because usually one is interested in finding out how quickly an existing DNA sequence can erode.

We used this method to calculate the loss probability of any DNA sequence motif, and we excluded insertions and deletions from this calculation.

### 4.3 Probabilities of finding, gaining, and losing DNA features

Usually, a specific function is encoded in DNA by several DNA sequences. For example, translation stop is encoded by three codons (`TGA`, `TAG`, `TAA`). We use the term DNA features to mean a set of DNA sequences that are associated with the same function. For every such DNA feature set, there is a complementary set of non-features, that is DNA sequences that are not associated with the feature’s function. For example, the non-feature set of stop codons would be all the other 61 codons. The probability that a DNA feature exists, is the sum of probabilities of every DNA sequence in that set (Section 4.2.1).

The probability that a DNA feature is gained via mutations, is the sum of probabilities of every non-feature sequence mutating to any feature sequence. If *F* denotes the feature set, and *µ*_*y→x*_ denotes the probability of a DNA sequence *y* mutating to a DNA sequence *x* (see Section 4.2.2), then:

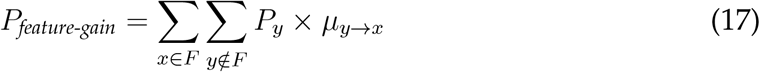

The probability that a DNA feature is lost via mutations is a conditional probability that given a feature exists, it mutates to any of the non-feature sequences.

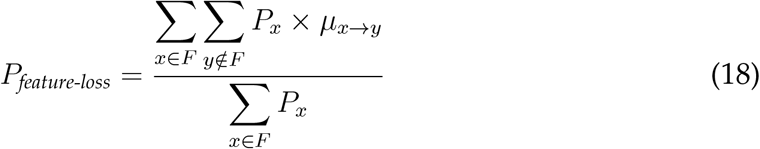

Because a feature set usually has many DNA sequences, a mutation can change a feature sequence such that the resulting sequence is also a part of the feature set. Thus we defined the probability (*P*_*feature-stay*_) that a feature does not erode because of mutations. Specifically, it is the sum of two probabilities. First is the probability that no mutation occurs in the DNA sequence (*P*_0_), and the second probability describes the event where the mutated sequence remains a part of the feature set.

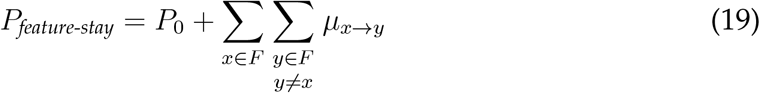

The probability that no mutation occurs (*P*_0_) in a DNA sequence of length *k* is described by Poisson distribution.

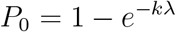

Because the mutation rate is biased (Table 1), the probability that no mutation occurs in a DNA sequence depends on its composition.

The probability that an `A` or a `T` mutates (*λ*_AT_), is thus described as:

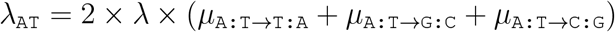

Likewise, the probability that a `G` or a `C` mutates (*λ*_GC_) is:

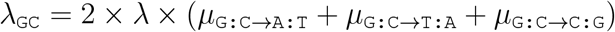

(Note that the general mutation rate, *λ*, is an average of *λ*_AT_ and *λ*_GC_.)

Thus the probability that a sequence of length *k*, containing *m* number of `A` and `T`, does not mutate is:

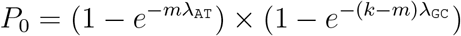

### 4.4 Probabilities of finding, gaining, and losing an ORF

#### 4.4.1 Probability of finding an ORF

A reading frame is a nucleotide sequence with a length that is a multiple of three. A reading frame that begins with a start codon (`ATG`), and ends with one of the three stop codons (`TAG`, `TGA`, `TAA`) is an open reading frame (ORF). This necessarily means that there are no stop codons within the sequence. Thus the probability of finding an ORF containing *k* codons including start and stop codons (*P*_*ORF*_) is:

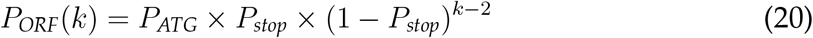

Here, *P*_*ATG*_ and *P*_*stop*_ are the probabilities of finding a start codon, and a stop codon by chance, respectively.

#### 4.4.2 Probability of gaining an ORF

As we defined in the previous section, an ORF has three requirements (start codon, stop codon, and no premature stop codon in the sequence). Thus an ORF can emerge due to mutations via three mechanisms. In each of these mechanisms, one requirement is initially absent whereas the other two are present. Then mutations cause the missing requirement to emerge while not disrupting the other two requirements. More specifically, the ORF can be gained via the following three mechanisms:

1. Gain of a start codon (*P*_*ATG-gain*_) while a stop codon continues to exist at the end of a reading frame (*P*_*stop-stay*_), and there is no emergence of stop codon within the reading frame (1 *− P*_*stop*_ *− P*_*stop-gain*_).
2. Gain of a stop codon (*P*_*stop-gain*_), while a start codon continues to exist at the beginning of a reading frame (*P*_*ATG-stay*_), and there is no emergence of stop codon within the reading frame.
3. Loss of a premature stop codon, at any of the *k* − 2 codon positions within the reading frame (*P*_*stop-gain*_). At the same time start and stop codons remain undisrupted by mutations, and no stop codon emerges at any of the other *k* − 3 positions.

Thus we define the probability of ORF gain (*P*_*ORF-gain*_) as:

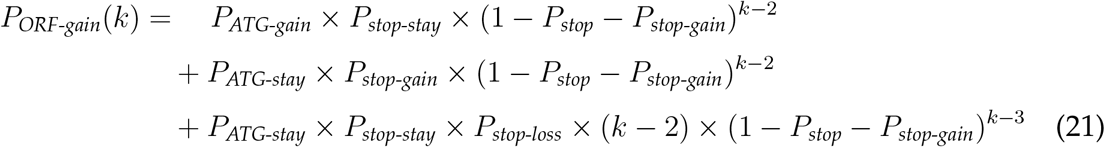

#### 4.4.3 Probability of ORF loss

ORF can be lost when any of its three requirements are lost. We thus define the conditional probability of ORF loss as:

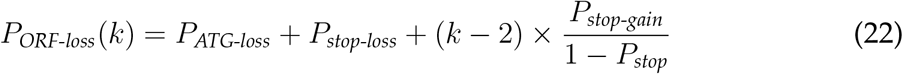

The last term in this equation describes the conditional probability of stop-gain, given the assumption that no stop codon exists within the ORF.

#### 4.4.4 Probability that ORF remains intact

We assumed that an ORF of a certain length remains intact if none of the necessary features are lost. However, the ORF sequence can mutate to cause non-synonymous changes in the translated protein sequence. This condition applies to all the three ORF probabilities described above. We define the probability that an ORF remains intact (*P*_*ORF-stay*_) as:

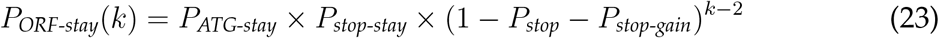

### 4.5 Probabilities of finding, gaining, and losing transcription

To calculate the probabilities of finding, gaining, and losing transcription, we first defined the minimum requirements for transcription. These requirements are a promoter, either a TATA-box or an Initiator (Inr) element, and a polyadenylation signal. Each of these three DNA sequence motifs consists of more than one DNA sequence.

#### 4.5.1 Definition of TATA-box, Inr and poly-A signal

##### TATA box

The consensus sequence of TATA-box is TATA [T/A] A [T/A] [G/A] (Basehoar *et al*., 2004) such that, any of the nucleotides enclosed within the square brackets, can exist in that specific position in the sequence. Another study used X-ray crystallography to identify mutations that are tolerated in TATA-box consensus sequence (Patikoglou *et al*., 1999). Thus there are eight consensus sequences, and at seven positions, a non-consensus nucleotide is tolerated (Table 2). This makes a TATA-box feature Position Accepted nucleotides set that contains 104 sequences. We defined the corresponding non-feature set by generating all 8-mer DNA sequences that do not belong to the TATA-box feature set.

**Table 2:**
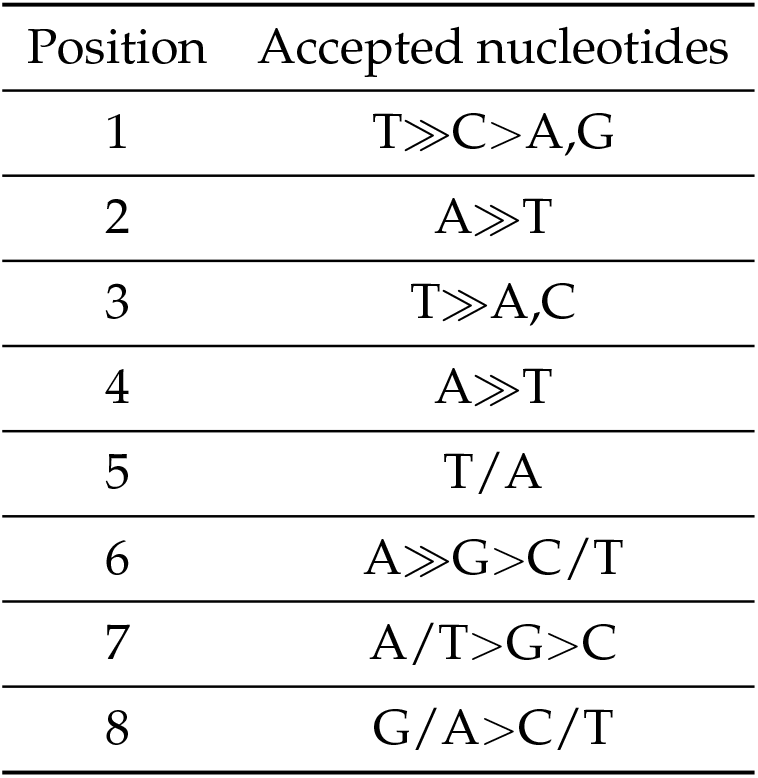
Nucleotide residues that are tolerated at different positions in a TATA-box (based on Patikoglou et *al*., 1999)

| Position | Accepted nucleotides |
| --- | --- |
| 1 | T>>C>A,G |
| 2 | A>>T |
| 3 | T>>A,C |
| 4 | A>>T |
| 5 | T/A |
| 6 | A>>G>C/T |
| 7 | A/T>G>C |
| 8 | G/A>C/T |

##### Inr

The consensus sequence of Inr is [T/G/C] [T/G/C] CA [T/G/C] [T/A], which gives 54 sequences in the Inr feature set (Kugel and Goodrich, 2017). We generated the corresponding non-feature set with all DNA 6-mers that are distinct from the 54 Inr sequences.

##### poly-A signal

There are four known poly-A signal variants – `AATAAA`, `ATTAAA`, `AGTAAA`, and `TATAAA` that compose the poly-A signal feature set (Proudfoot, 2011). The non-feature set consists of all other DNA 6-mers.

We calculated probabilities of finding, gaining and losing these features as we described in the previous section.

#### 4.5.2 Probability of transcription

For a a DNA region of a length, to be transcribed it should have a promoter the beginning of the DNA region, and a poly-A signal at the end of it. The transcribed DNA region should also not have any poly-A signal sequences within it except the one at the end of it. A transcribed DNA of length *l* nucleotides, contains *l −* 5 6-mers. Each of these 6-mers can be a poly-A signal with a probability, *P*_*polyA*_. Probability that none of these 6-mers is a poly-A signal is (1 *− P*_*polyA*_)^*l−*5^. Since we focus on protein coding genes in this work, we describe the length of transcript in terms of the length of protein it can encode. Assuming that untranslated regions have no effect on protein synthesis, the length of a transcript in number of nucleotides should be at least three times the length of the encoded protein in number of amino acids. Thus the probability that a DNA region produces a transcript that can harbor an ORF with *k* codons (*P*_*RNA*_) is:

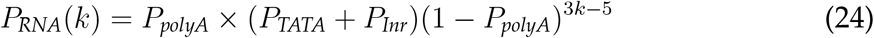

#### 4.5.3 Probability of gaining transcription

As we mentioned in the previous section, three requirements need to be met to produce a transcript of any given length. First, a promoter needs to be present. Second, a poly-A signal needs to be present at the end of the DNA region to be transcribed, and third, no poly-A signal should exist within this DNA region. Transcription can be gained if one of two of these requirements are already met and the third required feature emerges due to mutations. At the same time mutations do not destroy the existing features. It is possible that two or all the required features are missing and they emerge due to mutations. However such an event is highly improbable. Thus transcription gain can occur via three mechanisms.

In the first mechanism:

- A promoter (TATA-box or Inr) exists at the beginning of a DNA region and remains intact (*P*_*TATA-stay*_, *P*_*Inr-stay*_)
- No poly-A signal sequences are present within the DNA region and none emerges (*P*_*nopolyA-stay*_ = 1 *− P*_*polyA*_ *− P*_*polyA-gain*_)
- A poly-A signal emerges at the end of the DNA region (*P*_*polyA-gain*_) In the second mechanism:
- A poly-A signal is present at the end of the DNA region remains intact
- No poly-A signal sequences are present within the DNA region and none emerges
- A promoter emerges at the beginning of the DNA region (*P*_*TATA-gain*_, *P*_*Inr-gain*_) Finally, in the third mechanism:
- A promoter exists at the beginning of a DNA region and remains intact
- A poly-A site is present at the end of the DNA region remains intact
- One poly-A site that was present in the DNA region, is lost. (*P*_*polyA-loss*_)
- At every other site in the DNA region, no poly-A signal exists and none emerges

The total gain probability (*P*_*RNA-gain*_) of a transcript that can harbor an ORF with *k* codons would be the sum of the probabilities of the three gain mechanisms:

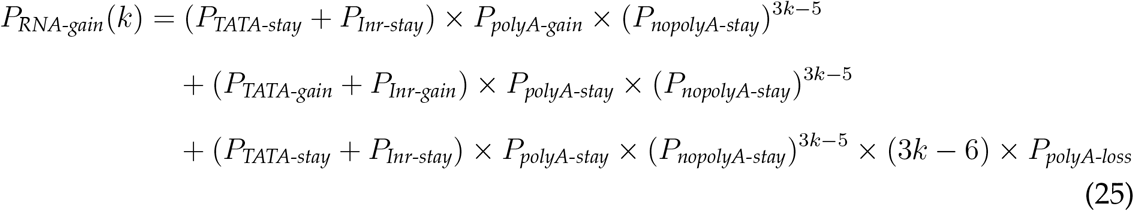

It is also possible that both the promoter and the poly-A signal were initially absent, and emerge via mutations. However, this probability is very small and is negligible.

#### 4.5.4 Probability of losing transcription

Transcription is lost when either the promoter is lost or when the poly-A signal is lost. Since two kinds of promoters can initiate transcription, there can be three kinds of transcription units. First that contains only a TATA-box, second that contains only an Inr, and the third that contains both these promoters. All these three transcription units must also have a poly-A signal. To calculate the conditional probability of promoter loss, we calculated the fraction of transcription units (*Q*) that belong to each of these three classes. For example, the fraction of transcription units that are TATA-only (*Q*_*TATA-only*_) is:

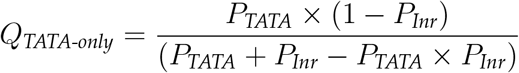

Here, the numerator denotes probability of finding a TATA-box but not an Inr, and the denominator denotes the probability of finding a promoter that can be a TATA-box, Inr, or both. Likewise, the fraction of Inr-only (*Q*_*Inr-only*_) and TATA-Inr promoters (*Q*_*both*_) are given by:

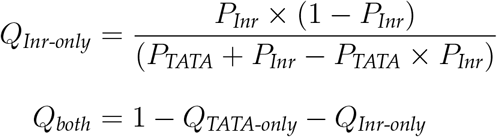

For a TATA-only transcription unit to lose a promoter, it should lose the TATA-box but simultaneously not gain an Inr. Likewise, an Inr-only transcription unit will lose a promoter when it loses the Inr but not gain a TATA-box. A transcription unit that has both the promoters must lose both of them. For all the three kinds of transcription units, transcription would be lost if poly-A signal is lost.

The conditional probability of transcription loss (*P*_*RNA-loss*_), given that transcription exists, is thus defined as:

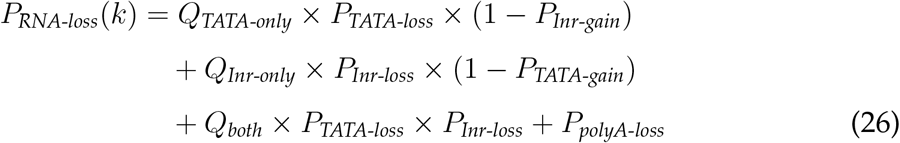

#### 4.5.5 Probability that transcription remains intact

Transcription remains intact if neither the promoter, nor the poly-A signal is lost. More specifically, we define this probability (*P*_*RNA-stay*_) as:

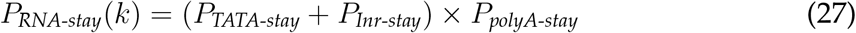

### 4.6 Interdependence of transcription and ORF probabilities

All the four poly-A signal variants contain a `TAA` in their sequence (position 3 – 5), which is also a stop codon. Since stop codons cannot exist inside an ORF, presence of an ORF reduces the possible number of poly-A signal sites in a DNA region. Specifically, a poly-A signal can exist in any of the three reading frames but it can overlap with the ORF only in one frame. Thus, out of 3*k* − 5 positions for a 6-mer (length of a poly-A signal) in an ORF with *k* codons, *k* positions cannot harbor a poly-A signal (Equation 24). Furthermore a poly-A signal cannot overlap with the start codon, thus reducing two more positions from all sites where a poly-A signal can exist (one position is already counted – if a poly-A signal overlaps with the first codon in the second frame, then the second codon becomes a stop codon). Overall, the possible number poly-A sites on an ORF with *k* codons is 2*k* − 5. Thus for a gene to emerge via gain of transcription, the DNA should not contain poly-A sites at these 2*k* − 5 positions. As a consequence, the probability of transcription gain for a specific transcript length, is higher when the DNA region contains an ORF than when it does not (Equation 25). We have ignored the probability of poly-A signal overlapping with a stop codon because it makes a very small difference in the probability of RNA gain. This is so because at least one stop codon (`TAA`) allows a poly-A signal to overlap with it in three frames.

Conversely to the effect of an existing ORF on transcription gain, stop codons are less probable in an existing transcript that does not contain any poly-A signal sequences within its sequence, than in an untranscribed DNA region. Specifically, the probability that a 6-mer contains a stop codon in the third position is same as the probability of finding a stop codon (*P*_*stop*_). If poly-A signals are excluded from these 6-mers, then the probability of finding a stop codon is *P*_*stop*_ *− P*_*poly-A*_. Conversely, lack of poly-A signals will increase the probability of amino acid coding codons and thereby their probability of mutating into a stop codon. Furthermore, the likelihood that an ORF is undisrupted my premature nonsense mutations, is lower when the DNA is already transcribed. Specifically:

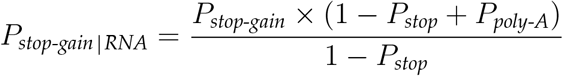

Overall the evolutionary dynamics of transcription and ORF gain, are not independent of each other. That is, presence of one feature makes the gain of the other more likely. The dependence becomes more prominent with increasing number of codons in the ORF of the proto-gene.

### 4.7 Probability of amino acid substitutions

We calculated the probability of amino acid substitutions based on nucleotide substitution in the corresponding codons. In this case, multiple codons that code for the same amino acid constitute a feature set (Section 4.3). Similarly when analysing hydrobhobic to non-hydrophobic substitutions, all codons that represent hydrophobic amino acids form a feature set (and *vice versa*).

## 5 Data availability

We performed all calculations using Julia programming language, and all scripts are freely available on GitHub: BharatRaviIyengar/DeNovoEvolution. Specifically, our model is implemented in the script `DeNovoProb.jl` which in turn uses the script `nucleotidefuncts.jl` for some basic functions.

## References

Acevedo, J. M., Hoermann, B., Schlimbach, T., and Teleman, A. A. 2018. Changes in global translation elongation or initiation rates shape the proteome via the Kozak sequence. Scientific Reports, 8(1): 4018.

Basehoar, A. D., Zanton, S. J., and Pugh, B. 2004. Identification and Distinct Regulation of Yeast TATA Box-Containing Genes. Cell, 116(5): 699–709.

Behrens, S. and Vingron, M. 2010. Studying the evolution of promoter sequences: A waiting time problem. Journal of Computational Biology, 17(12): 1591–1606.

Berg, J. M., Tymoczko, J. L., and Stryer, L. 2002. Biochemistry. W.H. Freeman, New York.

Blevins, W. R., Ruiz-Orera, J., Messeguer, X., and others 2021. Uncovering de novo gene birth in yeast using deep transcriptomics. Nature Communications, 12(1): 604.

Box, G. 1979. Robustness in the strategy of scientific model building. In Robustness in Statistics, pages 201–236. Academic Press.

Bucciantini, M., Giannoni, E., Chiti, F., and others 2002. Inherent toxicity of aggregates implies a common mechanism for protein misfolding diseases. Nature, 416(6880): 507–511.

Cano, A. V., Rozhoňová, H., Stoltzfus, A., McCandlish, D. M., and Payne, J. L. 2022. Mutation bias shapes the spectrum of adaptive substitutions. Proceedings of the National Academy of Sciences, 119(7): e2119720119.

Carvunis, A.-R., Rolland, T., Wapinski, I., and others 2012. Proto-genes and de novo gene birth. Nature, 487(7407): 370–374.

Choe, Y.-J., Park, S.-H., Hassemer, T., and others 2016. Failure of RQC machinery causes protein aggregation and proteotoxic stress. Nature, 531(7593): 191–195.

Dayhoff, M., Schwartz, R., and Orcutt, B. 1978. Model of evolutionary change in proteins. In Atlas of protein sequence and structure, volume 5, pages 345–352. National Biomedical Research Foundation, Silver Spring MD.

Dill, K. A. 1985. Theory for the folding and stability of globular proteins. Biochemistry, 24(6): 1501–1509.

Gardini, S., Cheli, S., Baroni, S., and others 2016. On nature’s strategy for assigning genetic code multiplicity. PLOS ONE, 11(2): e0148174.

Gerstein, M. B., Bruce, C., Rozowsky, J. S., and others 2007. What is a gene, post-ENCODE? History and updated definition. Genome Research, 17(6): 669–681.

Gonnet, G. H., Cohen, M. A., and Benner, S. A. 1992. Exhaustive matching of the entire protein sequence database. Science, 256(5062): 1443–1445.

Gramates, L. S., Agapite, J., Attrill, H., and others 2022. FlyBase: a guided tour of highlighted features. Genetics, 220(4).

Haberle, V. and Stark, A. 2018. Eukaryotic core promoters and the functional basis of transcription initiation. Nature Reviews Molecular Cell Biology, 19(10): 621–637.

Harris, K. and Nielsen, R. 2014. Error-prone polymerase activity causes multinucleotide mutations in humans. Genome Research, 24(9): 1445–1454.

Hartl, F. U. 2017. Protein misfolding diseases. Annual Review of Biochemistry, 86(1): 21–26.

Henikoff, S. and Henikoff, J. G. 1992. Amino acid substitution matrices from protein blocks. Proceedings of the National Academy of Sciences, 89(22): 10915–10919.

Hershberg, R. and Petrov, D. A. 2010. Evidence That Mutation Is Universally Biased towards AT in Bacteria. PLOS Genetics, 6(9): 1–13.

Hochberg, G. K. A., Liu, Y., Marklund, E. G., and others 2020. A hydrophobic ratchet entrenches molecular complexes. Nature, 588(7838): 503–508.

Iyengar, B. R., Choudhary, A., Sarangdhar, M., and others 2014. Non-coding RNA interact to regulate neuronal development and function. Frontiers in Cellular Neuroscience, 8: 47.

Jones, D. T., Taylor, W. R., and Thornton, J. M. 1992. The rapid generation of mutation data matrices from protein sequences. Bioinformatics, 8(3): 275–282.

Keeling, D. M., Garza, P., Nartey, C. M., and Carvunis, A.-R. 2019. Philosophy of Biology: The meanings of ‘function’ in biology and the problematic case of de novo gene emergence. eLife, 8: e47014.

Kim, Y., Sidney, J., Pinilla, C., Sette, A., and Peters, B. 2009. Derivation of an amino acid similarity matrix for peptide:MHC binding and its application as a bayesian prior. BMC Bioinformatics, 10(1).

Kosiol, C., Holmes, I., and Goldman, N. 2007. An empirical codon model for protein sequence evolution. Molecular Biology and Evolution, 24(7): 1464–1479.

Kozak, M. 1986. Point mutations define a sequence flanking the AUG initiator codon that modulates translation by eukaryotic ribosomes. Cell, 44(2): 283–292.

Kugel, J. F. and Goodrich, J. A. 2017. Finding the start site: redefining the human initiator element. Genes & Development, 31(1): 1–2.

Kyte, J. and Doolittle, R. F. 1982. A simple method for displaying the hydropathic character of a protein. Journal of Molecular Biology, 157(1): 105–132.

Le, S. Q. and Gascuel, O. 2008. An improved general amino acid replacement matrix. Molecular Biology and Evolution, 25(7): 1307–1320.

Lenhard, B., Sandelin, A., and Carninci, P. 2012. Metazoan promoters: emerging characteristics and insights into transcriptional regulation. Nature Reviews Genetics, 13(4): 233–245.

Long, M., Betrán, E., Thornton, K., and Wang, W. 2003. The origin of new genes: glimpses from the young and old. Nature Reviews Genetics, 4(11): 865–875.

Majic, P. and Payne, J. L. 2020. Enhancers facilitate the birth of de novo genes and gene integration into regulatory networks. Molecular Biology and Evolution, 37(4): 1165–1178.

Merchant, S. S., Prochnik, S. E., Vallon, O., and others 2007. The Chlamydomonas Genome Reveals the Evolution of Key Animal and Plant Functions. Science, 318(5848): 245–250.

Monroe, J. G., Srikant, T., Carbonell-Bejerano, P., and others 2022. Mutation bias reflects natural selection in Arabidopsis thaliana. Nature, 602(7895): 101–105.

Näsvall, J., Sun, L., Roth, J. R., and Andersson, D. I. 2012. Real-time evolution of new genes by innovation, amplification, and divergence. Science, 338(6105): 384–387.

Noderer, W. L., Flockhart, R. J., Bhaduri, A., and others 2014. Quantitative analysis of mammalian translation initiation sites by FACS-seq. Molecular Systems Biology, 10(8): 748.

Ohta, T. and Kimura, M. 1971. Amino acid composition of proteins as a product of molecular evolution. Science, 174(4005): 150–153.

Omotajo, D., Tate, T., Cho, H., and Choudhary, M. 2015. Distribution and diversity of ribosome binding sites in prokaryotic genomes. BMC Genomics, 16(1): 604.

Patikoglou, G. A., Kim, J. L., Sun, L., and others 1999. TATA element recognition by the TATA box-binding protein has been conserved throughout evolution. Genes & Development, 13(24): 3217–3230.

Prabh, N. and Rödelsperger, C. 2019. De Novo, divergence, and mixed origin contribute to the emergence of orphan genes in Pristionchus nematodes. G3 Genes|Genomes|Genetics, 9(7): 2277–2286.

Proudfoot, N. J. 2011. Ending the message: poly(A) signals then and now. Genes & Development, 25(17): 1770–1782.

Richard, P. and Manley, J. L. 2009. Transcription termination by nuclear RNA polymerases. Genes & Development, 23(11): 1247–1269.

Santangelo, T. J. and Artsimovitch, I. 2011. Termination and antitermination: RNA polymerase runs a stop sign. Nature Reviews Microbiology, 9(5): 319–329.

Schmid, M. and Jensen, T. H. 2018. Controlling nuclear RNA levels. Nature Reviews Genetics, 19(8): 518–529.

Schmitz, J. and Bornberg-Bauer, E. 2017. Fact or fiction: updates on how protein-coding genes might emerge de novo from previously non-coding DNA. F1000Research, 6(57).

Schneider, A., Cannarozzi, G. M., and Gonnet, G. H. 2005. Empirical codon substitution matrix. BMC Bioinformatics, 6(1).

Schrider, D. R., Houle, D., Lynch, M., and Hahn, M. W. 2013. Rates and Genomic Consequences of Spontaneous Mutational Events in Drosophila melanogaster. Genetics, 194(4): 937–954.

Shen, S., Kai, B., Ruan, J., and others 2006. Probabilistic analysis of the frequencies of amino acid pairs within characterized protein sequences. Physica A: Statistical Mechanics and its Applications, 370(2): 651–662.

Statello, L., Guo, C.-J., Chen, L.-L., and Huarte, M. 2021. Gene regulation by long non-coding RNAs and its biological functions. Nature Reviews Molecular Cell Biology, 22(2): 96–118.

Stewart, M. 2019. Polyadenylation and nuclear export of mRNAs. Journal of Biological Chemistry, 294(9): 2977–2987.

Tautz, D. and Domazet-Lošo, T. 2011. The evolutionary origin of orphan genes. Nature Reviews Genetics, 12(10): 692–702.

Vakirlis, N., Hebert, A. S., Opulente, D. A., and others 2017. A molecular portrait of de novo genes in yeasts. Molecular Biology and Evolution, 35(3): 631–645.

Van Oss, S. B. and Carvunis, A.-R. 2019. De novo gene birth. PLOS Genetics, 15(5): 1–23.

Whelan, S. and Goldman, N. 2001. A general empirical model of protein evolution derived from multiple protein families using a maximum-likelihood approach. Molecular Biology and Evolution, 18(5): 691–699.

Wimley, W. C. and White, S. H. 1996. Experimentally determined hydrophobicity scale for proteins at membrane interfaces. Nature Structural & Molecular Biology, 3(10): 842–848.

Wood, V., Gwilliam, R., Rajandream, M.-A., and others 2002. The genome sequence of Schizosaccharomyces pombe. Nature, 415(6874): 871–880.

Zhang, Z. and Gerstein, M. 2003. Patterns of nucleotide substitution, insertion and deletion in the human genome inferred from pseudogenes. Nucleic Acids Research, 31(18): 5338–5348.

Zhao, L., Saelao, P., Jones, C. D., and Begun, D. J. 2014. Origin and spread of de Novo genes in Drosophila melanogaster populations. Science, 343(6172): 769–772.

